# Computational design and evaluation of optimal bait sets for scalable proximity proteomics

**DOI:** 10.1101/2024.10.03.616533

**Authors:** Vesal Kasmaeifar, Saya Sedighi, Anne-Claude Gingras, Kieran R. Campbell

## Abstract

The spatial organization of proteins in eukaryotic cells can be explored by identifying nearby proteins using proximity-dependent biotinylation approaches like BioID. BioID defines the localization of thousands of endogenous proteins in human cells when used on hundreds of bait proteins. However, this high bait number restricts the approach’s usage and gives these datasets limited scalability for context-dependent spatial profiling. To make subcellular proteome mapping across different cell types and conditions more practical and cost-effective, we developed a comprehensive benchmarking platform and multiple metrics to assess how well a given bait subset can reproduce an original BioID dataset. We also introduce GENBAIT, which uses a genetic algorithm to optimize bait subset selection, to derive bait subsets predicted to retain the structure and coverage of two large BioID datasets using less than a third of the original baits. This flexible solution is poised to improve the intelligent selection of baits for contextual studies.

## Introduction

Spatial partitioning is crucial to the function of biological systems, which contain multiple layers of complexity, from organs to cells down to the smallest subcellular entities and molecules^1,2^. Eukaryotic cells separate diverse biochemical activities into distinct subcellular components, such as membrane-bound and membraneless organelles. This separation allows proteins to operate in specific environments, and improper localization often leads to disease. Studying protein localization under different conditions is essential to understanding their subcellular functions and links to disease^3,4^.

Various techniques have been used to investigate the subcellular localization of proteins. Currently, fluorescence-based imaging and biochemical fractionation coupled with mass spectrometry (MS) are most common^3,5–8^. While useful for studying most cellular components, fluorescence-based imaging requires assessing the localization of each protein individually, a critical bottleneck for whole-proteome studies with multiple conditions^3^. Biochemical fractionation and MS is more scalable and has been used to delineate the proteomes of numerous cellular components, and the overall subcellular organization^4–6,9^. However, some cellular structures are challenging to fractionate, including most membraneless organelles^10^.

To mitigate these issues, proximity labelling was developed^11^. It involves fusing an enzyme to a protein of interest (*i.e*., the bait) and expressing it in relevant cellular contexts. Subsequent addition of the enzyme’s substrate results in the covalent labelling of adjacent proteins (*i.e*., preys) in living cells (**Fig. 1a**). In BioID^12^, a modified biotin ligase produces an activated, reactive biotinoyl-AMP cloud that covalently biotinylates the lysine ε-amines of proteins within ∼10 nm^13^. The biotinylated proteins are then purified using streptavidin, and identified by MS. Protein identifications are scored against negative controls (*e.g*., using significance analysis of interactome (SAINT)^14–16^) to identify high-confidence proximal interactions and generate spatially resolved proteomic data. Additionally, while each bait identifies its own set of proximal proteins, multiple baits generate rich data matrices that can be analyzed using prey-centric approaches, such as correlation analysis or non-negative matrix factorization (NMF)^17^, to identify preys with similar interaction profiles that are likely to colocalize. These data can then be used to reconstruct cellular (or subcellular) organization^4,18^. NMF is a linear dimensionality reduction technique that decomposes a bait-prey matrix into a basis matrix and a score matrix. It is well suited for defining organellar composition because it soft-assigns prey proteins into a pre-specified number of components simultaneously— reflecting the cellular reality, in which many proteins have no singly defined subcellular localization. While the user is free to set the number of components, a heuristic in g:Profiler^19^ based on Gene Ontology Cellular Component (GO:CC) analysis of the preys assigned to each component is commonly used^4,18^. Components identities are annotated following a guilt-by-association principle, using GO terms or similar reference sets.

**Figure 1:**
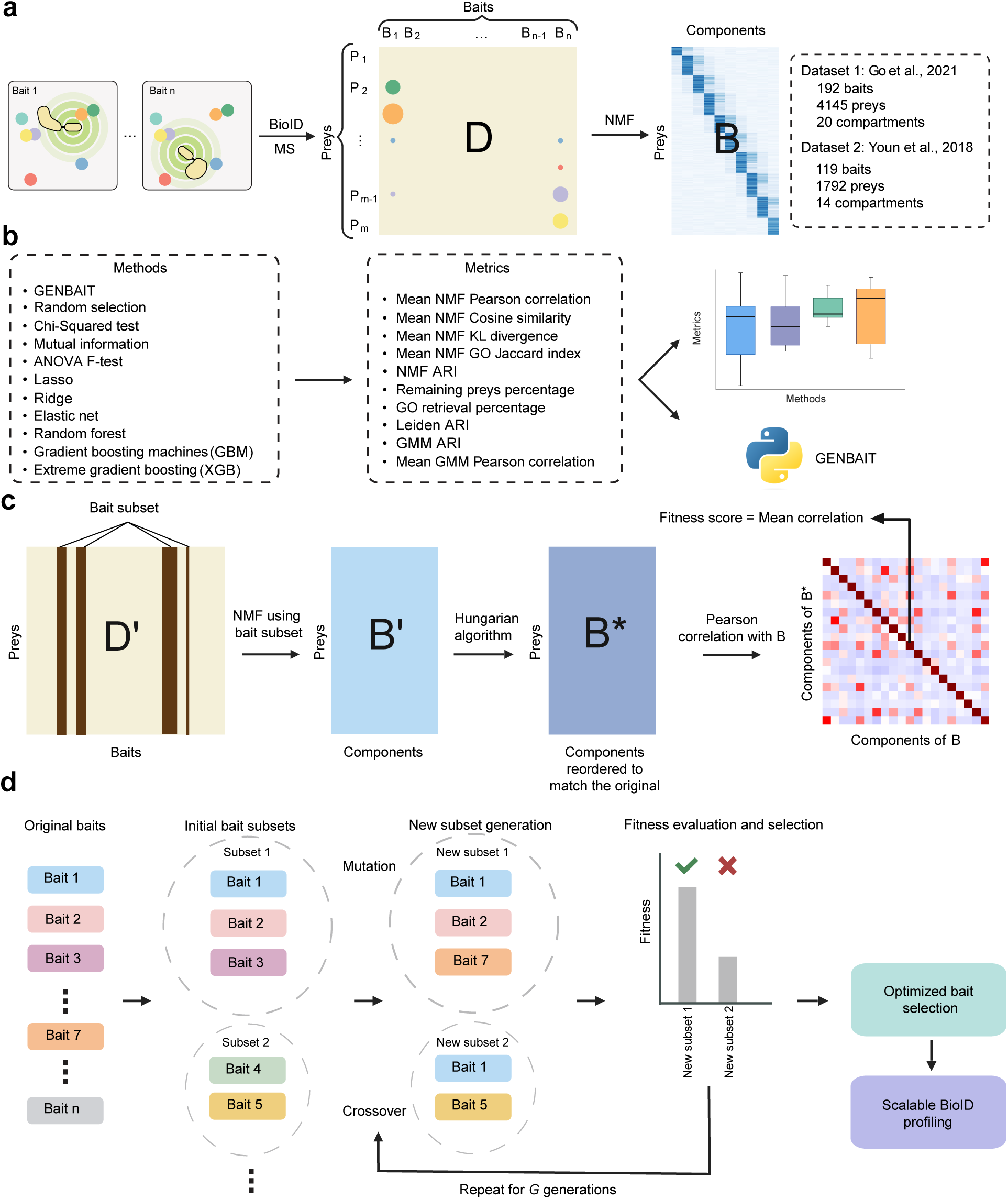
GENBAIT workflow and evaluation. **a.** Proximal labeling data are acquired by MS. Interaction scoring is performed to generate a matrix of baits and high-confidence preys (D). NMF is used to soft-assign preys into a predefined number of components (B) based on GO:CC terms. **b.** We compared GENBAIT’s performance to nine feature selection methods and random selection (as a baseline) using 10 different metrics. **c.** Schematic of the evaluation procedure (fitness function). Bait selection generates a subset (D’) of the original dataset (D). NMF is used to sort the subset’s preys into components (B’), and the Hungarian algorithm is used to create a matrix (B*) in which the components of B’ are aligned with those in the full dataset (B). Then, Pearson correlations between corresponding components are calculated. The mean of the diagonal values is used as the fitness score in the genetic algorithm. **d.** Workflow of the genetic algorithm for optimizing BioID bait subsets. Randomly selected initial bait subsets undergo mutation and crossover operations, followed by fitness evaluation. High-scoring subsets are used in the next generation to iteratively define the optimal subset for scalable BioID profiling.

This method has been used to generate BioID-based maps of the human cell (192 baits)^4^, cytosolic mRNA-associated granules and bodies (119 baits)^18^, and nuclear bodies (140 baits)^20^. However, these studies were performed in a single cell line under consistent growth conditions, largely because of the costs associated with profiling large numbers of baits. A means to select bait subsets that can recapitulate the structures of these interactomes with similar coverage would enable subcellular protein localization mapping in different contexts. From a machine learning perspective, this can be framed as a feature selection problem^21^, in which a subset of features (*i.e*., baits) is selected that maintains the strong predictive capacity of the full set. While there are a multitude of algorithms for feature selection that have been extensively applied to problems in biological data analysis^22,23^, none have been evaluated for their ability to generate bait subsets for scalable BioID profiling studies using formalized metrics, and no bespoke approaches have been proposed.

Here, we develop GENBAIT, a genetic algorithm-based strategy for BioID bait subset selection and introduce a novel benchmarking platform to quantify the quality of the BioID bait subsets, which we use to compare GENBAIT’s performance with that of nine existing statistical and machine learning-based feature selection algorithms. While all selection methods performed markedly better than random subset selection, GENBAIT outperformed the other methods across several metrics. Additionally, we formalize a set of recommendations to help researchers choose the optimal method with which to derive bait subsets from existing BioID datasets for scalable subcellular profiling. A Python package implementing both GENBAIT and the 10 metrics for assessing bait subset quality is available at https://github.com/camlab-bioml/genbait.

## Results

### A computational platform to design and evaluate BioID bait subsets

To evaluate the quality of a given subset of baits returned by a feature selection method *in silico*, we employed two existing published BioID datasets, each acquired in HEK293 cells in standard culture conditions: (1) the Human Cell Map V.1 (humancellmap.org), which used 192 baits to quantify the intracellular localizations of 4,145 high-confidence preys across the entire cell^4^, and (2) a more focused map of cytosolic mRNA-associated granules and bodies, which includes 119 baits and 1792 high-confidence preys^18^ (**Fig. 1a**). NMF optimization in the original publications defined 20 and 14 components, respectively, for these datasets. Then, we developed a benchmarking resource comprising 10 different evaluation metrics that quantified how well a given subset of baits captured the localizations determined in the original dataset and applied this to both BioID datasets individually (**Fig. 1b**).

Next, we developed a computational method to select bait subsets across multiple BioID experiments that can reproduce the interaction network of the full bait/data set, enabling scalable profiling. Given their success at solving similar discrete optimization problems^24–28^, we adopted a genetic algorithm-based search strategy informed by the typical BioID data analysis workflow. Implementing this strategy requires specifying a *fitness function* that quantifies how well a subset conserved the subcellular localizations observed with the full bait set. We developed a fitness function that mimics the iterative data analysis process^4,18^ to interpret subcellular colocalizations from BioID data (**Fig. 1c**). Specifically, for a proposed bait set, we applied NMF to the bait-prey matrix, retaining only the proposed bait subsets with the same number of components as the original NMF fit. We then computed each component’s Pearson correlation between the original and subset NMF components and used the Hungarian algorithm^29^ to re-order the components to best match the original. Finally, a single fitness score was computed by taking the mean of the correlations of the best-matched components between the original and subset (**Methods**).

Starting with an initial set of randomly generated bait subsets, the algorithm iteratively applied crossover (combining parts of two or more subsets to create new ones), and mutation (randomly changing baits in the subsets to introduce variability) operations^30^ to generate new subsets, then evaluated each with a fitness function. The subsets with the highest fitness scores were selected to generate subsequent subsets, and the operations were repeated. Over successive generations, this process generated optimized bait subsets for scalable BioID profiling (**Fig. 1d**). We refer to this algorithm as GENBAIT. An open-source Python package implementing GENBAIT as well as the metrics to assess bait subset quality is available at https://github.com/camlab-bioml/genbait_reproducibility.

### Contrasting bait subset selection by GENBAIT to random selection

To compare the efficacy of a GENBAIT-determined bait subset to a randomly selected subset of the same size, we intuitively selected ∼1/3 of the baits in each dataset (60 and 40 for datasets 1 and 2, respectively), as a reasonable example. We then recomputed their NMF representations, then measured their correlations with the NMF results of the full datasets (**Fig. 2a**). While these random subsets captured most components (as shown by the high correlation coefficients on the heatmap diagonals), they missed important components in both datasets. For dataset 1, no subset NMF components mapped to original components 2, 4, 11 and 18, representing the nucleolus, nucleoplasm, endosome, and chromatin, respectively. Similarly, for dataset 2, no subset components mapped to original components 3, 10, 12, 13, representing the nucleolus, spliceosome, nucleus, and cytoskeleton, respectively. Notably, when GENBAIT was used to generate optimized bait subsets, all original components were present in the subset NMF, and the correlation coefficients, which indicate how well each component was represented in each dataset, were increased (**Fig. 2b**).

**Figure 2:**
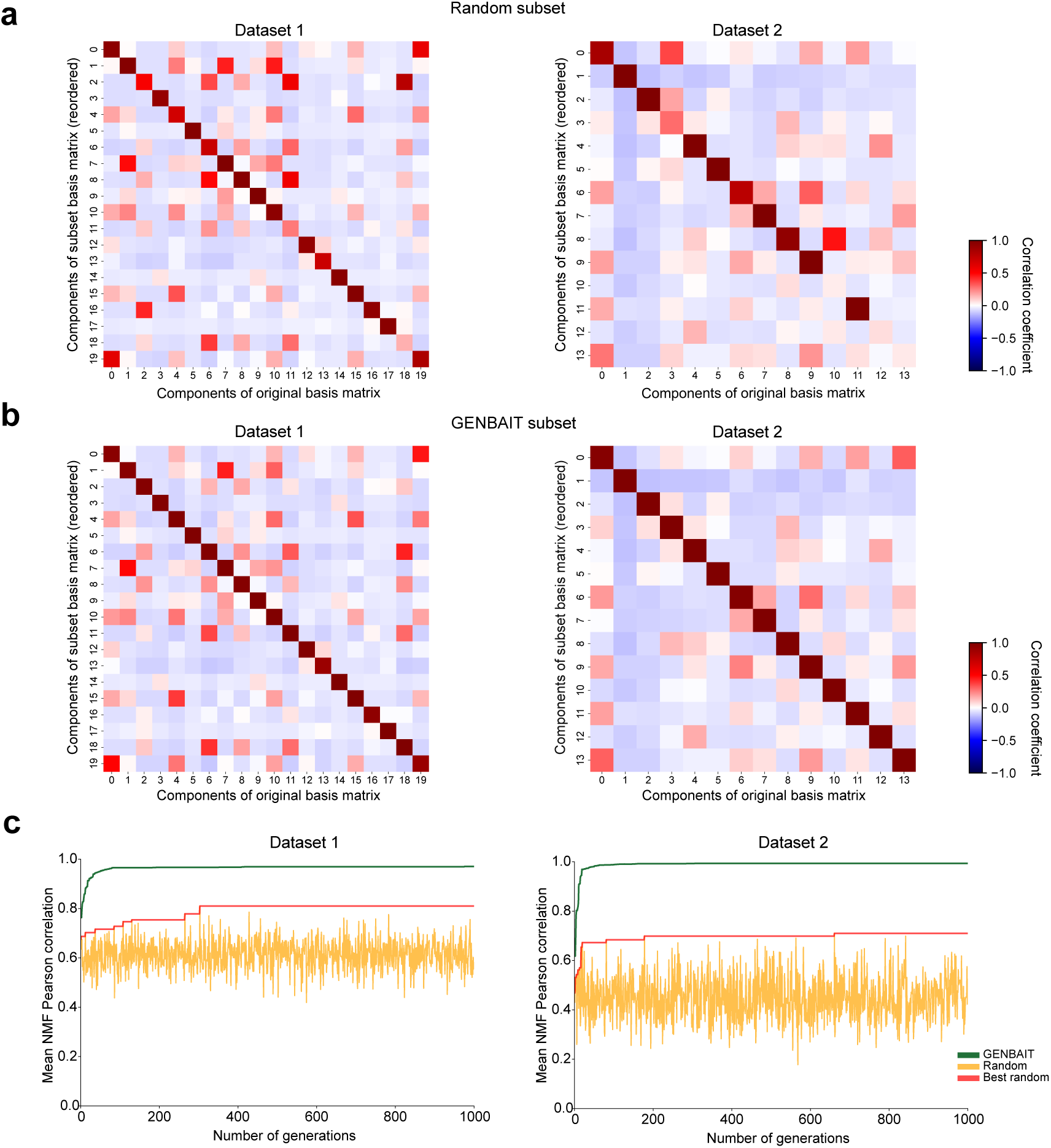
Comparing bait selection with GENBAIT against random selection. **a–b.** Correlation heatmaps between the original NMF results from datasets 1 and 2 and randomly selected subsets of each. Diagonal values indicate correlations between corresponding components of the original and subset basis matrices. **c.** Comparison of mean NMF Pearson correlation scores over 1000 generations of bait subsets using either GENBAIT or random selection or best random subset.

As GENBAIT created many bait subsets over multiple generations (1000 in our workflow), we next asked if random selection could generate a subset as high-scoring as a GENBAIT-determined subset if given the same computational resources (quantified using the mean diagonal NMF scores of its components). Therefore, we generated a random subset for each GENBAIT iteration and compared their fitness scores. The best random subset, reflecting the highest score achieved across all random selections during the entire process, is also shown. Notably, GENBAIT subsets quickly approached a maximum mean NMF Pearson correlation score (within < 100 generations), while the randomly selected subsets and best random subset did not, even after 1000 generations (**Fig. 2c**).

We also compared GENBAIT to an alternate heuristic approach to bait subset selection, in which each bait was ranked according to the number of preys it captured, and the top *N* baits were retrieved for a bait subset target size of *N* (**Fig. S1)**. In dataset 1, no components in the subset NMF mapped to original components 5, 12, 13, 14, and 17, representing the endoplasmic reticulum, actin cytoskeleton, microtubule cytoskeleton, mitochondrial inner membrane, and cytoplasmic stress granules, respectively. In dataset 2, components 6 and 10, representing cytoplasmic stress granules and the spliceosome, respectively, were present in the subset NMF, but their correlations with the original components were lower than with GENBAIT, indicating that GENBAIT better preserved the NMF components of the original datasets.

### Comprehensive benchmarking across multiple feature selection methods and metrics

We next used our benchmarking pipeline to quantify the performances of GENBAIT and 10 other feature selection methods (**Fig. 1b**), including statistical tests of the associations between baits and components (*e.g.*, analysis of variance (ANOVA), mutual information and Chi-squared)^31^, variants of sparse regression that select baits with non-zero coefficients for the subset (lasso, ridge and elasticnet)^32,33^, and tests that measure the feature importance per bait (random forest, gradient boosting machines (GBM)^34,35^ and extreme gradient boosting (XGB)^36^). All methods were compared using subsets containing 30–80 baits, and 10 random seeds over a range of components (15-25 for dataset 1 and 9-19 for dataset 2).

We contrasted these methods across 10 metrics that measure how their bait subset’s NMF results reflected those of the full datasets. These included metrics to explicitly quantify the similarity between the original and subset bait NMF results, using the Pearson correlation, cosine similarity, and Kullback–Leibler (KL) divergence. We also computed the Jaccard index of enriched GO terms and the adjusted Rand index (ARI) between the original and subset scores following hard assignment of each prey to a component (**Fig. 3a–e**, **Supplementary Table 1**). This revealed several characteristics of the performance of bait subset algorithms. First, as expected, all methods markedly outperformed random subset selection with both datasets and all evaluation metrics. Secondly, GENBAIT moderately outperformed the other feature selection methods across all these NMF-based metrics for both datasets. Interestingly, the broad trends indicated that after GENBAIT, the machine learning-based feature importance methods (random forest, gradient boosting machine (GBM), and extreme gradient boosting (XGB)) performed best, followed by the linear penalty methods (lasso, ridge, and elastic net regression), then the statistical methods (ANOVA *F*, Mutual info and Chi-squared). These trends were broadly consistent over various numbers of NMF components (**Fig. S2**).

**Figure 3:**
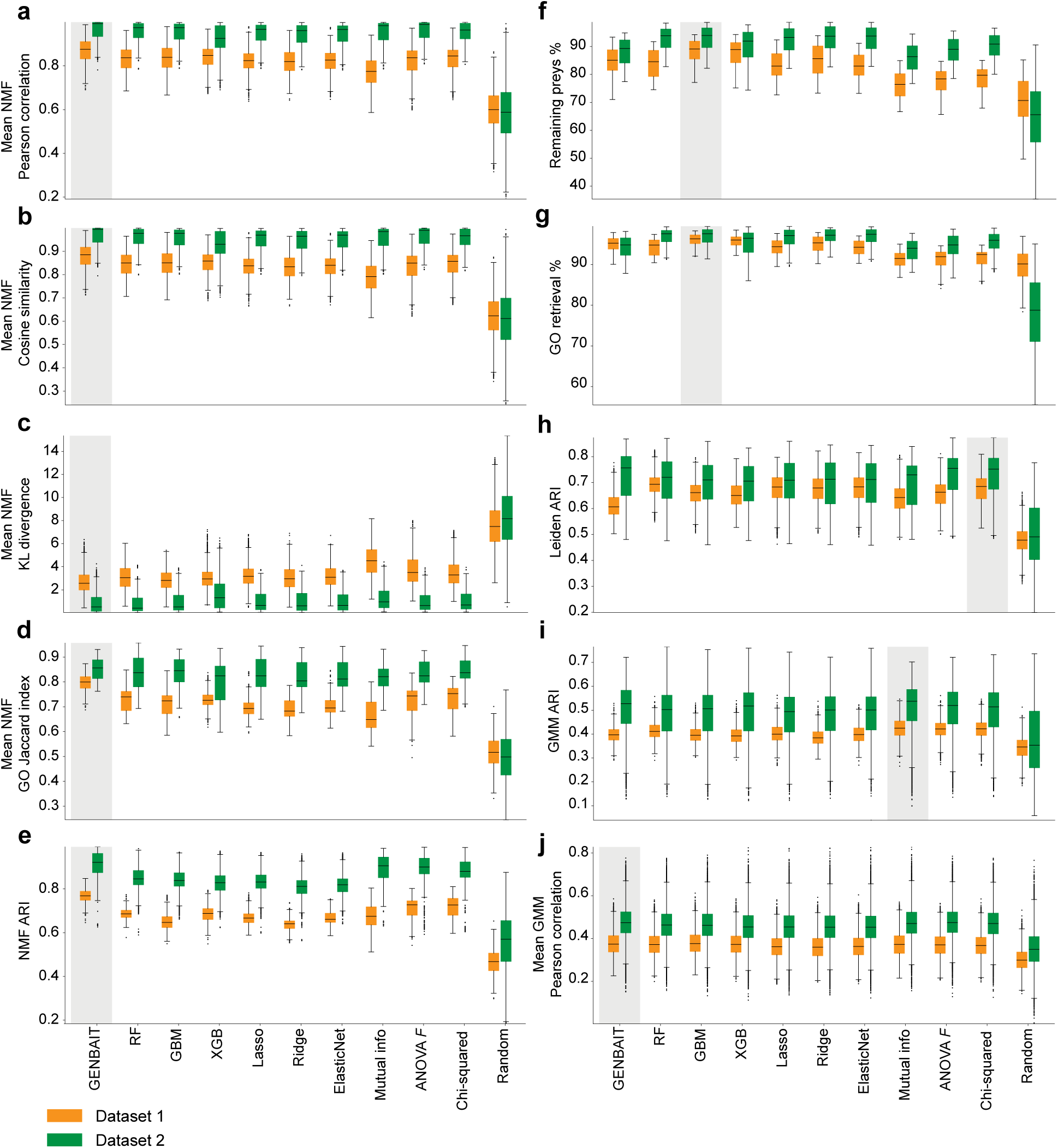
Benchmarking GENBAIT and other methods using NMF- and non NMF-derived metrics. **a–j.** Boxplots comparing GENBAIT, benchmarking methods, and random selection based on mean NMF Pearson’s correlation scores (**a**); NMF-based metrics (mean cosine similarity (**b**), KL divergence (**c**), GO term Jaccard index (**d**), and ARI (**e**)); and non-NMF metrics, including the percentages of remaining preys (**f**) and GO retrievals (**g**); Leiden clustering of KNN graphs at three resolutions (the ARI between common preys across 10 random seeds are shown; **h**); and GMM hard clustering (**i**) and soft clustering (**j**) at four numbers (dataset 1: 15, 20, 25, 30; dataset 2: 5, 10, 15, 20). Each method selected 30–80 baits from a range of components (15-25 for dataset 1 and 9-19 for dataset 2) over 10 random seeds. Box plots show median values; hinges are the 25th and 75th percentiles; whiskers indicate 1.5× the interquartile range (IQR). Highest median values are shaded in grey.

To understand the properties conserved by their bait subsets, we also contrasted the methods using a range of NMF-independent metrics, including the percentage of original preys still confidently assigned to baits in the subset, the overlap in annotated GO terms between the original and subset matrices, and the consistency of their clustering using different algorithms. As above, all feature selection methods significantly outperformed random bait selection across all evaluation metrics (**Fig. 3f–j**, **Supplementary Table 2**). Interestingly, the GBM bait subsets retrieved the highest percentages of the original preys and had the highest GO term similarity between the original and subset components (**Fig. 3f–g**). In contrast, statistical feature selection methods provided optimal cluster overlap, with the mutual information and chi-squared methods having the highest ARI values upon Leiden^37^ and Gaussian mixture model (GMM)^38^ hard clustering (**Fig. 3h–i**). Finally, GENBAIT ranked highest for contrasting cluster mean correlations between the original and subset bait matrices using GMM soft clustering (**Fig. 3j**). These trends were generally conserved with varying numbers of clusters (**Fig. S3**).

Together, these results reveal that GENBAIT particularly excels at optimizing NMF-derived metrics and mean GMM correlations. This is especially relevant to our study, as it emphasizes the importance of soft clustering metrics like these to accurately capture proteins that localize to multiple components.

### Robustness of subcellular map reconstruction at different bait subset sizes and random seeds

We next used GENBAIT, random selection, and each alternative method to generate different subsets containing 30–80 baits. For each number of baits, we generated 10 different random subsets for each dataset to assess robustness to the initial bait set. This was necessary because of the inherent randomness in the initialization processes of the algorithms. In GENBAIT, this randomness comes from the initial bait subset, which is selected randomly. In other methods, the randomness arises from the random train-test split of data, which differs with each seed. We then measured the mean NMF Pearson correlation scores for these subsets. For dataset 1, GENBAIT generated high-quality subsets over the entire size range; however, performance dropped notably for subsets with < 40 baits (**Fig. 4a**). Variance in the mean NMF Pearson correlation scores was low, indicating the consistently high quality of the bait subsets GENBAIT generated. The same conclusions were drawn for dataset 2 (**Fig. 4b**), though the differences in the mean NMF Pearson correlation scores between GENBAIT and the alternative methods were much smaller. Interestingly, XGB’s performance markedly decreased with smaller subsets, despite a strong performance when averaged over all subset sizes (**Fig. 4b**). Because the variances in mean NMF Pearson correlation scores between GENBAIT and the alternative methods were so small for subsets with small subset size, we repeated our analysis of dataset 2 with subsets of 15–35 baits. In this analysis, the mean NMF Pearson correlation scores for GENBAIT were higher than those of competing methods for subsets with ≥ 18 baits; however, GENBAIT’s performance was slightly worse (< 10%) with smaller subsets (**Fig. 4c**).

**Figure 4:**
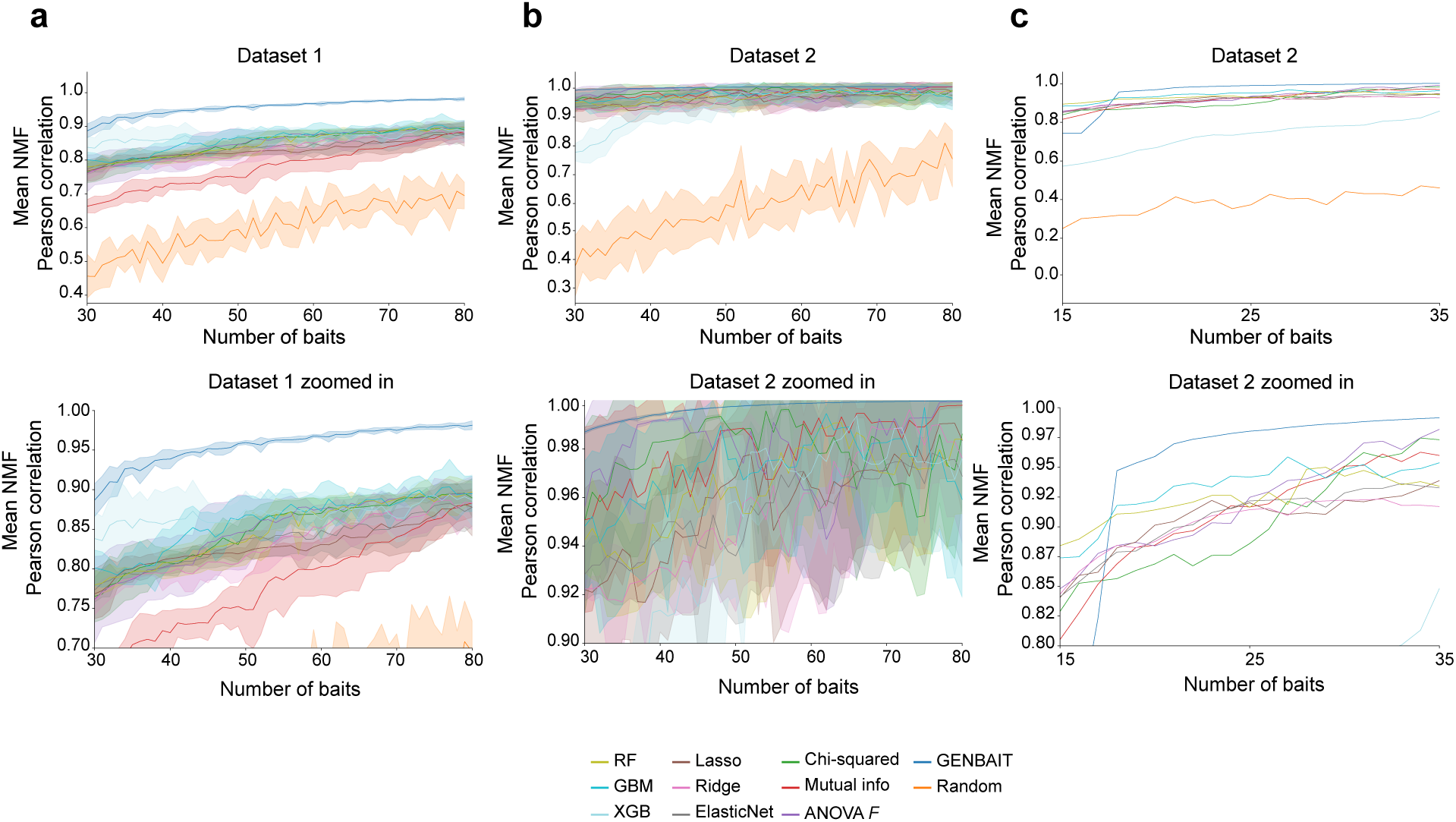
Impacts of varying the subset size and random initializing seed on the performance of bait subset selection methods. **a–c.** Mean NMF Pearson correlation scores are shown for each feature selection method across different bait subset sizes. Colored lines represent the mean NMF scores of each feature selection method, and shaded areas indicate the variability across different random seeds.

### Recommendations for bait subset selection for BioID

To facilitate a comparison of GENBAIT and alternative feature selection methods, we generated overall scores and ranked them (**Fig. 5**). We computed average scores for each evaluation metric and scaled them such that the minimum value was 0 and the maximum was 1, ensuring consistency across different metrics. The overall scores for each method were then determined by averaging these scaled scores across all metrics, except for the primary metric, which is the mean NMF Pearson correlation score. Finally, these overall scores were also scaled to a 0–1 range for consistency. GENBAIT ranked highest for both datasets, while the performance of other feature selection methods varied between them. Notably, while metrics based on clustering techniques favored methods like chi-squared, ANOVA F-test, and mutual information, these methods primarily focus on detecting the main localization of proteins. However, our objective is to preserve all localizations of a protein within the subsets, which may not be fully captured by these feature selection approaches. GENBAIT, on the other hand, can successfully retain the information for all localizations of a prey protein and, therefore, be a suitable choice for protein multilocalization studies.

**Figure 5:**
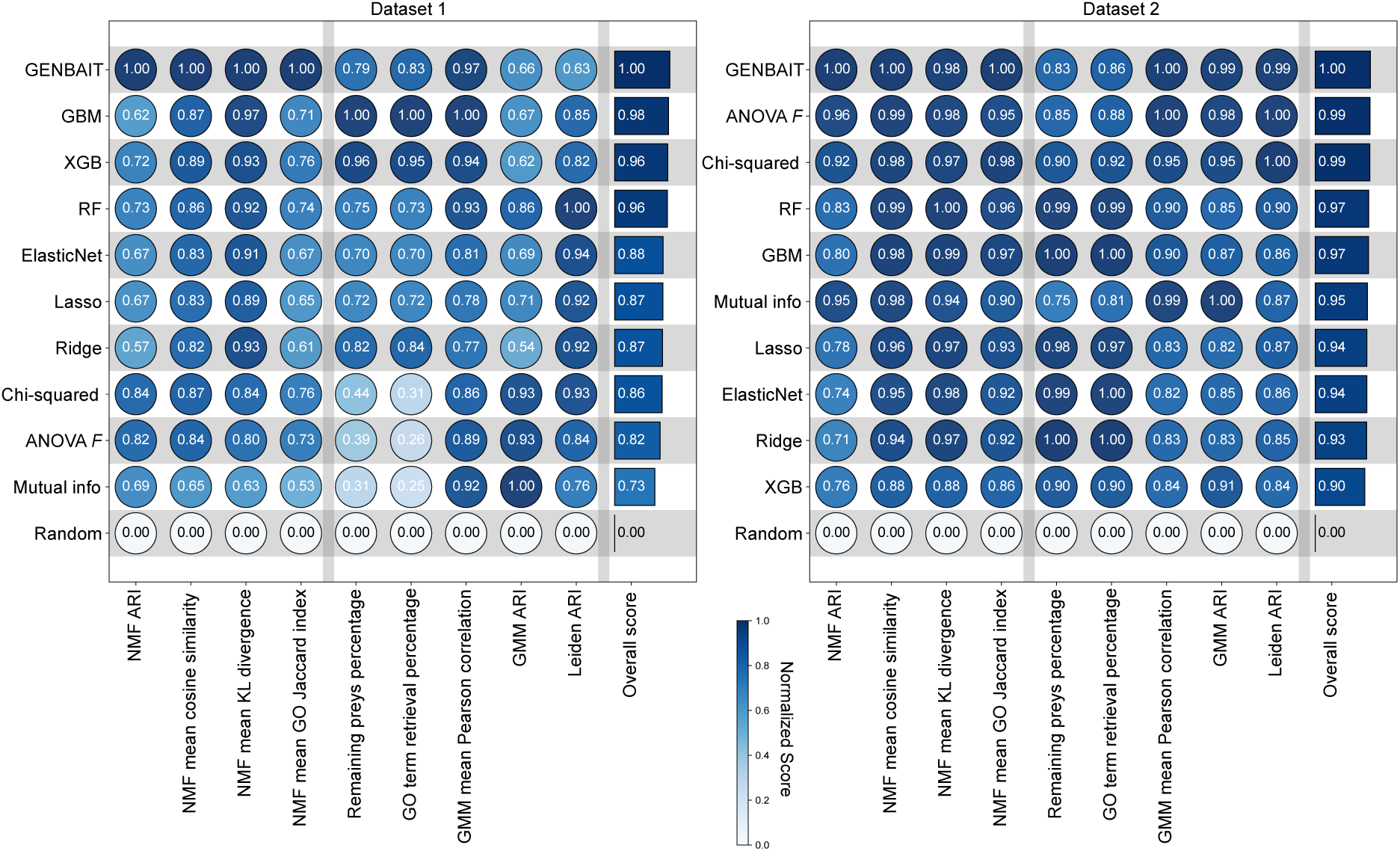
Scoring of bait subset selection methods by metric. A heatmap of the scores of various feature selection methods across multiple metrics. The color of each circle represents its normalized score. The overall scores are normalized average scores for each method over all metrics.

Overall, our results demonstrated that no single feature selection method universally outperformed the others on every metric. Therefore, we suggest that researchers prioritize feature selection methods over random subset selection, carefully selecting the optimal method using our benchmarking metrics. To guide this process, we provide a selection framework based on our results in **Fig. 6**. These guidelines are a starting point for subset selection and should be further tailored by the end user’s knowledge of the dataset and specific analytical objectives. Overall, the choice of method will hinge on the desired accuracy, available computational resources, and urgency. If the primary aim is to maintain high similarity between the cluster structures of the original and subset maps, we recommend GENBAIT, which consistently outperformed other methods across NMF-derived metrics and GMM soft clustering. However, its extensive optimization process requires more computational power. Our analysis suggests that a system with a minimum of 16 GB of RAM is needed to handle the intensive computations and large datasets. When reliable results are needed quickly, regression-based methods such as lasso, ridge, or elastic net are advisable. These methods are less computationally demanding than GENBAIT and performed well in our benchmarking pipeline, particularly in statistical association tests, they offer a practical compromise between speed and accuracy. When computational resources are constrained but robust results are required, ensemble methods like random forest, GBM, or XGB are recommended. These methods balance computational efficiency with the ability to uncover underlying patterns in the data. In our evaluations, they showed relatively good performance in both NMF-based metrics and in maximizing the numbers of retained preys and GO term retrievals, making them suitable for multilocalization analysis.

**Figure 6:**
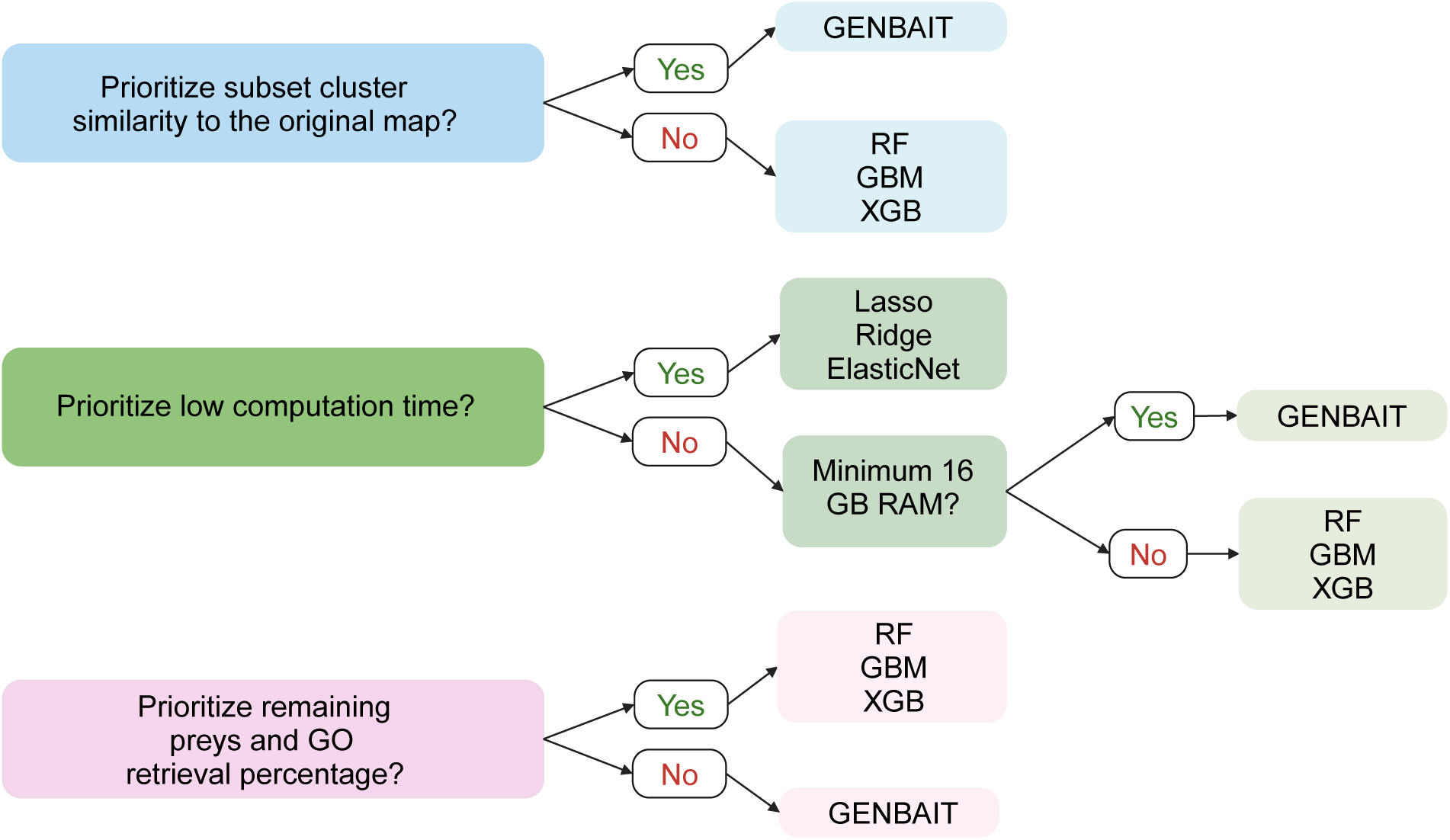
Guidelines for choosing a BioID bait subset selection method. The optimal method will depend on the question asked, the experimental priorities, and the available computational resources.

## Discussion

Our study demonstrates the potential of feature selection methods, including a novel genetic algorithm-based approach GENBAIT, in addressing the challenge of selecting optimal bait subsets for BioID experiments. By developing and applying a comprehensive evaluation pipeline with 10 diverse metrics, we have provided a robust framework for assessing and comparing these methods in the context of proximity proteomics research.

One of the key contributions of our work is the demonstration that established feature selection methods can be effectively adapted for BioID bait selection, offering a significant improvement over random selection. Additionally, our introduction of a novel set of benchmarking metrics allows researchers to make informed decisions when choosing the most appropriate method for their specific datasets and research goals. These metrics serve as practical tools that can guide researchers in selecting bait subsets that maintain the structural and functional integrity of the original dataset.

GENBAIT showed strong performance across several metrics, particularly those derived from NMF-based analyses and GMM soft clustering. However, it did not universally outperform all other methods across every metric. Methods based on machine learning and statistical approaches also performed well, particularly in metrics related to primary protein localization. This highlights the importance of selecting a feature selection method that aligns with the specific objectives of the study, rather than relying solely on one approach.

Despite the strengths of GENBAIT, several limitations should be acknowledged. First, although our computational approach provided valuable insights, we did not experimentally validate any of the bait subsets generated by GENBAIT or other methods. Additionally, our study did not include a comparison with a baseline established by a well-trained human expert, who might manually select bait subsets based on domain knowledge. Such a comparison could offer valuable context for understanding the strengths and weaknesses of algorithmic versus expert-driven selection.

Another consideration is the computational demand of GENBAIT. While not prohibitively time-consuming, GENBAIT’s thorough optimization process requires more computational resources than other methods evaluated. This could pose a challenge for researchers with limited computational infrastructure. Future iterations of GENBAIT could focus on improving computational efficiency without sacrificing the thoroughness that contributes to its strong performance.

In conclusion, our study highlights the utility of feature selection methods for BioID bait subset selection and provides a comprehensive framework for evaluating these methods. While GENBAIT offers a highly competitive approach, particularly for studies focused on multi-localization, it is not a one-size-fits-all solution. Researchers should carefully consider their specific research objectives, available resources, and the trade-offs between accuracy and computational efficiency when selecting a feature selection method. Future studies will focus on optimizing GENBAIT’s parameters to enhance computational efficiency and robustness. The next iteration will incorporate a lab-in-the-loop approach, integrating experimental validation with computational predictions to improve consistency and reliability.

## Supplementary information

### Supplementary Figures

**Supplementary Figure 1.**
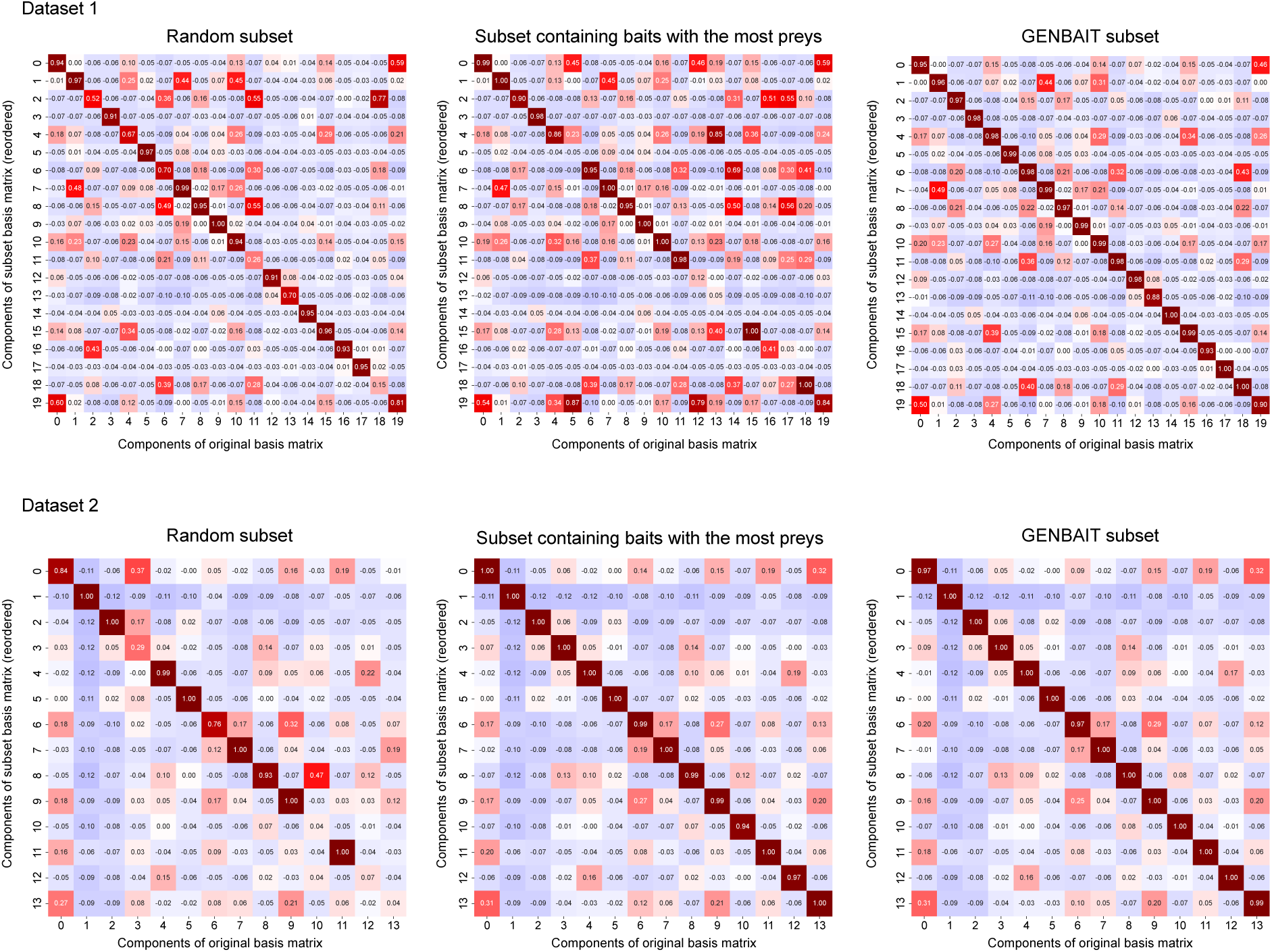
Correlations between the original and subset basis matrices. Heatmaps of the correlations between components of the original and subset basis matrices generated by different bait selection methods. Corresponding components are located on the diagonal.

**Supplementary Figure 2.**
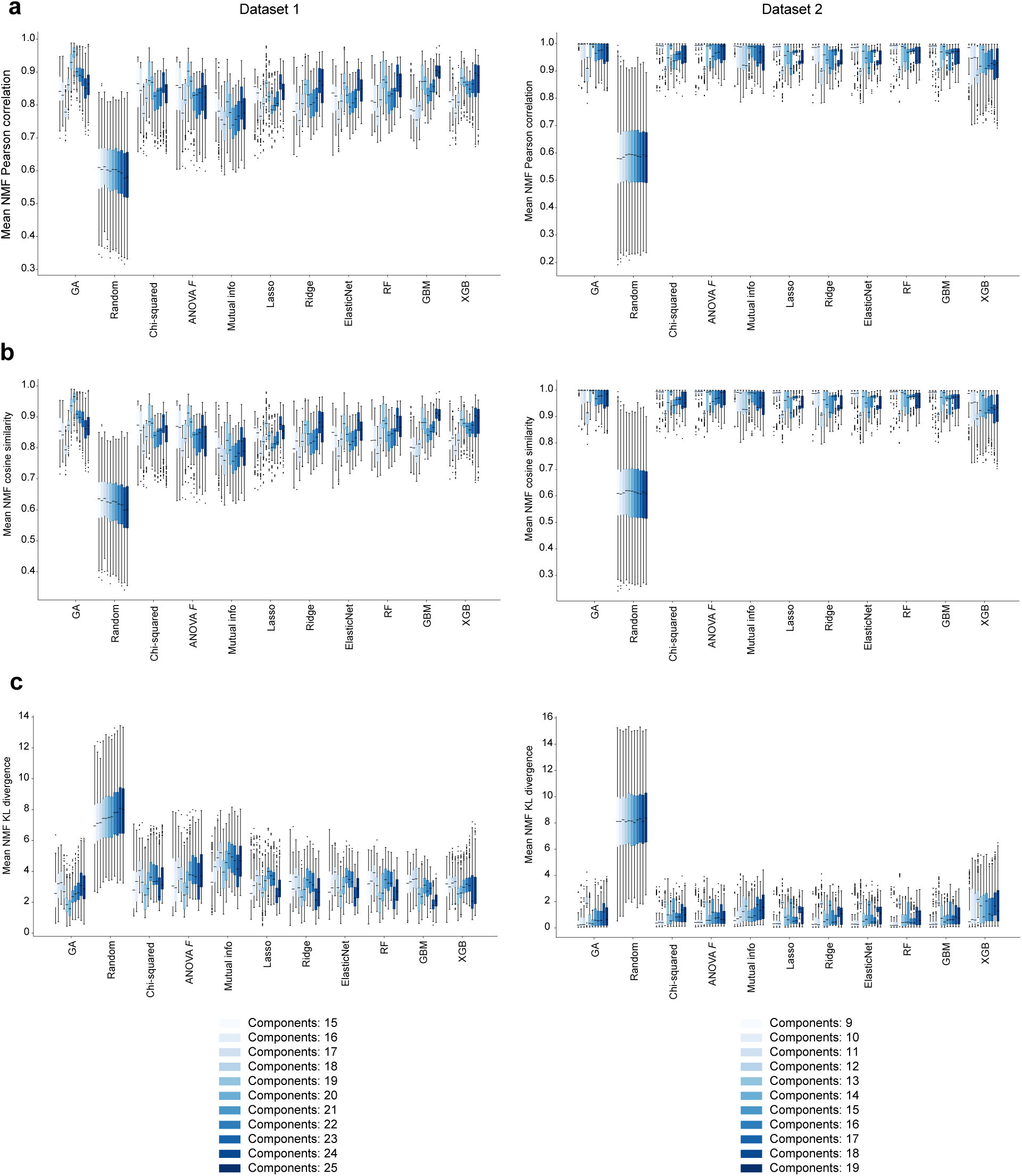
Benchmarking GENBAIT and other methods using NMF-derived metrics with various component numbers. **a–c.** For each feature selection method, mean NMF Pearson correlations, cosine similarities, and KL divergence scores were calculated using component numbers ranging from five less than in the original dataset to five more. Box plots show median values; the hinges are the 25th and 75th percentiles; whiskers indicate 1.5× the IQR.

**Supplementary Figure 3.**
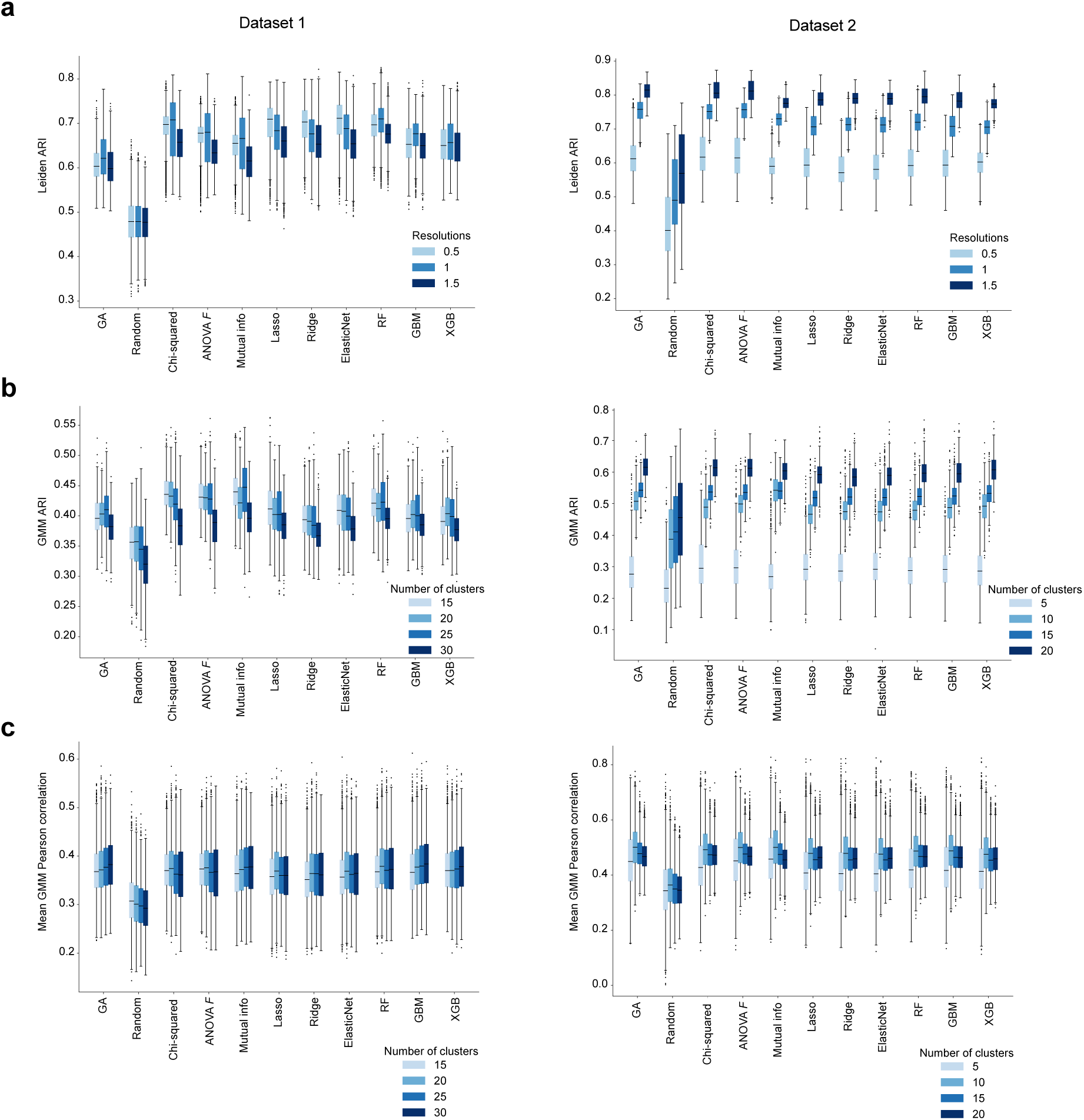
Benchmarking GENBAIT and other methods using non-NMF-derived metrics with various resolutions and cluster numbers. **a.** For each feature selection method, the Leiden ARI was calculated at the indicated resolutions. **b–c.** For each feature selection method, the GMM ARI (**b**) and mean GMM Pearson correlation (**c**) were calculated with the indicated numbers of clusters. Box plots show median values; hinges are the 25th and 75th percentiles; whiskers indicate 1.5× the IQR.

### Supplementary Table captions

**Supplementary Table 1. Average median values of different methods across NMF-derived metrics**

This table shows average median values of different bait subset selection methods across NMF-derived metrics.

**Supplementary Table 2. Average median values of different methods across non-NMF-derived metrics**

This table shows average median values of different bait subset selection methods across non-NMF-derived metrics.

## Methods

### Data preprocessing

The datasets were preprocessed using a custom pipeline to ensure data quality and consistency. First, the average spectral counts for each prey protein identified in the negative controls was subtracted from its average spectral counts observed for a given bait, giving a corrected average. Preys were then filtered to include only high-confidence interactions (*i.e*., those with a Bayesian false discovery rate^16^≤ 0.01). These data were pivoted to create a matrix with the baits in rows and the preys in columns, which was filled with corrected average spectral count values. The MinMaxScaler from the scikit-learn library^39,40^ was applied to scale the data, ensuring that feature values were normalized to fall within a given range on a per-feature basis. If a list of primary baits was specified, the matrix was further reduced to include only those baits and their corresponding non-zero preys. The preprocessing steps were encoded in Python, utilizing the Pandas^41^ library for data manipulation and the scikit-learn^42^ library for scaling.

### Benchmarking methods

#### Data preparation

Each normalized dataset was divided into two subsets: 80% for training and 20% for testing. This division was performed randomly, using a set seed to ensure reproducibility. To avoid data leakage, we used non-negative least squares (NNLS)^43^ fitting during the preparation process. NMF was first applied to each full dataset to decompose it into scores and basis matrices with the original number of components. The basis matrix components were then used to create the target variables for feature selection, by assigning each prey to the component with its highest component score. Subsequently, NMF was applied to the training subset to obtain its scores and basis matrices. For the testing subset, the basis matrix was initialized with zeros and calculated using NNLS fitting to ensure that the decomposition was consistent with that of the training subset. This approach helped maintain the integrity of the dataset by preventing information from the test set from leaking into the training process. The resulting matrices were combined to form the full set for training and testing. The final training and test sets were then created, where X_train and X_test represent the transposed data matrices, and y_train and y_test contain the target variables for the training and test subsets, respectively.

#### Feature selection methods

We used multiple feature selection methods to identify the most informative features:

- Chi-squared tests^42^ were used to evaluate the independence of each feature from the class label (assigned based on the component with highest value after NMF decomposition). The top features were selected based on their chi-squared scores.
- ANOVA *F*-Tests^44^ were used to identify features with significant differences across the classes. Features with the highest scores were selected.
- Mutual information^45^ was used to measure the dependency between each feature and the class label. Features with the highest scores were selected.
- Lasso regression^46^ was used to perform feature selection by imposing an L1 penalty, which drives some coefficients to zero, effectively performing feature selection. Features with the largest absolute coefficients were selected.
- Ridge regression^46^, which uses an L2 penalty to shrink coefficients, helps minimize the loss by adding a penalty equivalent to the square of the magnitude of coefficients. Features with the largest absolute coefficients were selected.
- Elastic net regression^47^ was used to select features by balancing the L1 and L2 penalties, thereby benefitting from the sparsity of lasso and the regularization of ridge. Features with the largest absolute coefficients were selected.
- Random forest^48^ was used to estimate feature importance based on how much each feature decreases the impurity of the splits. The features that contributed most to reducing the impurity were selected based on these importance scores.
- GBM^49^ was used to evaluate feature importance. Features that contributed the most to reducing the loss function over multiple boosting iterations were selected.
- XGB^50^ was used to assess feature importance based on its gradient boosting algorithm. The top features were selected based on their importance scores, focusing on those that contributed the most to reducing the loss function.

#### Iterative analysis across seeds

The feature selection process was iterated over 10 random seeds (from 0 to 9) to assess the consistency of the selected features. The results from each seed were aggregated and analyzed to understand the stability of each feature selection method.

### Random bait subset selection

To establish a baseline for comparison, we generated 1000 random subsets of features by randomly selecting a set of indices from the normalized dataset, with the subset size falling within a specified range between 30-80. We then evaluated their utility as a reference point against which the performance of more systematic feature selection methods could be assessed, using a modified version of the evalSubsetCorrelation function that was designed to assess the subset’s representativeness of the original dataset based on NMF component correlations. The fitness function generated the mean of the diagonal values of the correlation matrix, optionally applying a gradient penalty for subsets with negative correlations.

### Bait subset selection with GENBAIT

After data preprocessing, GENBAIT was employed to identify the most informative subset of baits from the datasets.

#### Algorithm initialization

An initial population of solutions was generated, with each individual represented as a binary vector. The length of the vector corresponded to the number of baits in the dataset, with each element indicating the presence (1) or absence (0) of a particular bait. A predetermined random seed was used to initialize the population, ensuring the reproducibility of the results.

#### Fitness function

The fitness of each individual was determined using a custom fitness function, *evalSubsetCorrelation*, which assessed the subset’s representativeness of the original dataset based on the correlations between their corresponding NMF components. Subsets outside the pre-specified size range were heavily penalized to enforce size constraints. For valid subsets, NMF was applied to extract basis matrices, followed by alignment using the Hungarian algorithm^29^ to ensure that the components were ordered as in the original dataset. The fitness score was calculated using the correlation matrix’s diagonal values, with penalties for negative correlations to discourage unrepresentative feature combinations.

The fitness function, *f*(*I*), for an individual *I* in the genetic algorithm is defined as follows:

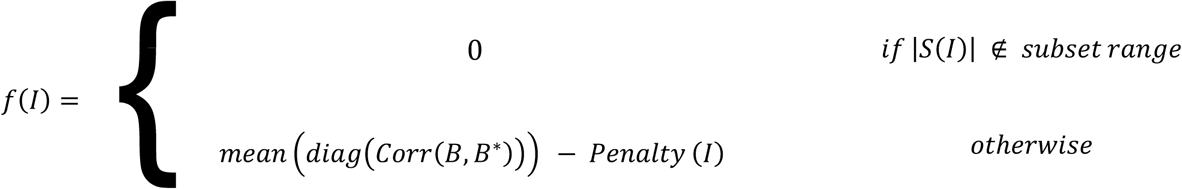

Where:

- |*S*(*I*)| is the size of the feature subset represented by the individual *I*.
- *subset range* is the allowable size range for the feature subset.
- *B* is the basis matrix from NMF applied to the original dataset.
- *B*^∗^is the reordered basis matrix from NMF applied to the subset of data corresponding to *I*.
- *Corr* (*A* − *B*) calculates the correlation matrix between matrices *A* and *B*.
- *diag* (*X*) extracts the diagonal elements of matrix *X*.
- *Penalty* (*I*) is a function that applies a penalty based on the number of negative values in the diagonal of the correlation matrix, calculated as:

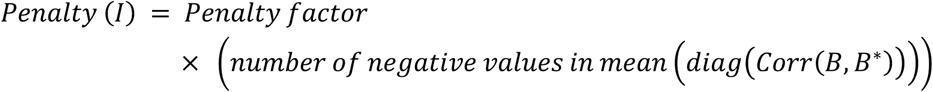
- *Penalty factor* is a predefined constant.

#### Genetic operators

Crossover and mutation were implemented to generate new solutions^51^. Crossover was performed using a two-point crossover method (tools.cxTwoPoint) with a specified probability (cxpb). Mutation involved flipping bits in the individual’s binary representation with a mutation probability (mutpb). Individuals were selected for the next generation based on fitness, using a tournament selection method (tools.selTournament).

#### Algorithm execution

GENBAIT ran for a defined number of generations (n_generations), each involving selection, crossover, and mutation. The algorithm’s progress and population dynamics were tracked using a logbook, and the best-performing individuals were recorded in a hall-of-fame (hof).

#### Computational environment and resources

The GENBAIT algorithm was implemented in Python, using the DEAP^52^, NumPy^53^, Pandas libraries for evolutionary computations, numerical operations, and data processing, respectively. NMF decomposition was performed using the scikit-learn library, and the Hungarian algorithm was applied through the SciPy^54^ library’s linear_sum_assignment function.

#### Algorithm parameters

Specific parameters used in GENBAIT, including population sizes, crossover and mutation probabilities, and the number of generations, were chosen based on preliminary experiments to balance computational efficiency and the quality of the feature selection process. GENBAIT outputted an optimized subset of baits for each dataset, with high representativeness and low redundancy, which were used in downstream analyses, including clustering and biological interpretation.

### Validation metrics

#### Mean NMF cosine similarity and KL divergence scores

Subsets generated by random selection, GENBAIT, or alternative methods were subjected to NMF, and their subset basis matrices were aligned with the original dataset as above. Instead of calculating the Pearson correlations between corresponding components, we calculated their cosine similarities and KL divergences across 10 random seeds.

#### NMF ARI and mean NMF GO Jaccard index

Subsets generated by random selection, GENBAIT, or alternative methods were subjected to NMF, and their subset basis matrices were aligned with the original dataset as above. To calculate the ARI, each prey was assigned to the component with the highest value. Using these assignments, we obtained ARI values between original data and subsets. The Adjusted Rand Index (ARI) measures the similarity between two clusterings by comparing the number of pairs of elements that are consistently clustered together or apart in both clusterings, adjusting for chance. The adjusted_rand_score function in sklearn.metrics.cluster was used to compute ARI values. A higher ARI indicates that the clustering structure in the subset closely matches that of the original dataset, with a value of 1 representing perfect agreement. We also used g:profiler to evaluate the biological relevance of the identified components. We extracted the names of the preys in each component identified by NMF in both the original and subset datasets and used g:profiler API to identify related GO terms for each component. We calculated the Jaccard index for corresponding components to quantify the similarity and consistency between the GO terms for the original and subset datasets, then averaged the indices for all clusters to obtain the mean Jaccard index. The Jaccard index measures the similarity between two sets by dividing the size of the intersection by the size of the union of the sets. In this context, a higher Jaccard index indicates greater overlap between the GO terms associated with the original and subset datasets, reflecting better consistency in the biological relevance of the clustering. This value provided a quantitative means to assess the biological relevance and accuracy of the clustering methods.

#### Graph construction using the K-nearest neighbors (KNN) algorithm

For each dataset, we generated a KNN graph using the kneighbors_graph function in the sklearn.neighbors module^55^. The number of neighbors (*k*) was set to 20, and the connectivity mode was used for graphing. The KNN graph was then converted into an igraph object, with vertices representing individual baits and edges indicating neighbor relationships.

#### Leiden clustering

Leiden clustering was applied to the KNN graph using the leidenalg library. The RBConfigurationVertexPartition method was used with a resolution parameter set to explore different levels of community detection^56^. The algorithm partitioned the graph into clusters, and the membership of each vertex (bait) was recorded. Clustering was performed at various resolutions (0.5, 1, 1.5) to assess the robustness of the cluster structures.

Clustering was performed separately for the original dataset and all generated subsets. For each subset, a KNN graph was constructed and clustered using the same parameters as for the original data. The adjusted_rand_score function in sklearn.metrics.cluster was used to compute ARI values between the cluster memberships in the original and subset datasets, providing a measure of the similarity in their clustering structures. The entire process was iterated over 10 random seeds (ranging from 0 to 9) to assess consistency. Results were aggregated and stored using pickle for subsequent analysis. The analysis was parallelized using the concurrent.futures.ProcessPoolExecutor to enhance computational efficiency.

ARI values and cluster counts for each resolution and method were analyzed to determine the degree of cluster preservation. A comparative analysis was conducted between GENBAIT, other machine learning methods, and random bait selection to evaluate GENBAIT’s effectiveness in maintaining biologically relevant clusters.

#### GMM hard clustering

For each bait subset, GMM was used to assign each feature to a cluster, using the GaussianMixture function from sklearn.mixture. GMMs with varying numbers of clusters were fitted to the data. ARI values were computed for the original dataset and each subset using adjusted_rand_score from sklearn.metrics. This measure helped evaluate how well the clustering structure of the subsets preserved that of the original dataset. The analysis was repeated across 10 random seeds (ranging from 0 to 9) to assess the consistency of the clustering results. The results were aggregated and analyzed to understand the stability of the clustering across different methods and cluster numbers.

#### GMM soft clustering

GMM was applied to assign probability distributions to the clusters for each feature, indicating its likelihood of belonging to each cluster. The predict_proba function from the GaussianMixture model was used to obtain these soft assignments. The average correlations between clusters in the original dataset and the subsets were computed by calculating the correlation matrix between the probability distributions of the original and subset datasets, then reordering the clusters to match. The mean of the diagonal values was used as a measure of the average correlation.

Like hard clustering, the soft clustering analysis was conducted across multiple random seeds. The results were compiled to compare the average cluster correlations across different cluster numbers and methods.

### GO term retrieval percentage

The GO analysis began by loading gene annotation file (GAF)^57,58^ data into a structured format. Each entry in the GAF file, adhering to the GAF 2.1 specification, was parsed to extract critical information including the database identifier, object symbol, and GO ID.

The core of the analysis hinged on comparing GO terms between the original dataset and subsets derived *via* different feature selection methods (or random selection). Each subset’s genes were mapped to GO:CC terms, with a focus on identifying and retaining terms within a specified maximum term size to mitigate the influence of broader terms focusing on a certain level of specificity. This mapping yielded a concise list of GO:CC terms for further analysis. The comparative analysis aimed to quantify the overlap in GO terms between the original dataset and the subsets. This was achieved by calculating the percentage of common GO terms, providing a measure of similarity and preservation of biological relevance across different feature selection methodologies. The analysis extended to evaluate the impact of varying the number of features selected, offering insights into the trade-offs between feature selection granularity and GO term conservation. To ensure the robustness and consistency of the results, the entire GO analysis process was iterated across 10 random seeds (ranging from 0 to 9). This approach enabled the assessment of stability and variability in GO term overlap and similarity metrics across different iterations, providing a comprehensive understanding of the methods’ performance.

#### Percentage of remaining preys

We next quantified how well the feature selection methods preserved relevant biological information in the original datasets. For each subset generated, we calculated the proportion of non-zero prey proteins to assess the retention of significant preys. This was accomplished by transforming normalized expression data into a NumPy array, isolating subsets, and counting non-zero elements across a predefined range of number of components (15-25 for dataset 1 and 9-19 for dataset 2) to capture different levels of feature stringency. The process was iterated to ensure the robustness of the findings, which were aggregated for subsequent visualization and analysis.

### Benchmarking the methods using different metrics and calculating overall scores

To compare GENBAIT’s performance with that of other feature selection methods, we calculated average scores for each evaluation metric, which were normalized to 0–1 to ensure the comparability of different metrics. Overall scores for each method were calculated by averaging the normalized scores across all metrics, excluding the main metric (the mean NMF Pearson correlation score). This approach allowed us to objectively assess and rank the performance of each method across multiple evaluation criteria.

## Data availability

SAINT file for dataset 1: https://humancellmap.org/resources/downloads/saint-latest.txt SAINT file for dataset 2: https://prohits-web.lunenfeld.ca/GIPR/supplementary_data.php?projectID=17&m_num=m3

GO Annotation File (GAF): https://geneontology.org/docs/go-annotation-file-gaf-format-2.1/

## Code availability

GENBAIT python package: https://github.com/camlab-bioml/genbait

GENBAIT reproducibility: https://github.com/camlab-bioml/genbait_reproducibility

## Acknowledgements

We thank High-Fidelity Science Communications for manuscript editing. We thank members of the Gingras and Campbell labs for helpful discussions.

## Authors and Affiliations

**Lunenfeld-Tanenbaum Research Institute, Mount Sinai Hospital, Sinai Health System, Toronto, Ontario, Canada**

Vesal Kasmaeifar, Saya Sedighi, Anne-Claude Gingras, Kieran R. Campbell

**Department of Molecular Genetics, University of Toronto, Toronto, Ontario, Canada**

Vesal Kasmaeifar, Saya Sedighi, Anne-Claude Gingras, Kieran R. Campbell

**Department of Computer Science, University of Toronto, Toronto, Ontario, Canada**

Kieran R. Campbell

**Department of Statistical Sciences, University of Toronto, Toronto, Ontario, Canada**

Kieran R. Campbell

**Ontario Institute for Cancer Research, Toronto, Ontario, Canada**

Kieran R. Campbell

**Vector Institute, Toronto, Ontario, Canada**

Kieran R. Campbell

## Contributions

Project conception: A.-C.G., V.K., K.R.C. Result interpretation and manuscript writing: V.K., K.R.C., A.-C.G.,. Data analysis and software development: V.K., K.R.C., Visualization: V.K., S.S.

## Corresponding authors

Correspondence to Anne-Claude Gingras and Kieran R. Campbell.

## Ethics declarations

Competing interests

The authors declare no competing interests.

